# Predicting coarse-grained representations of biogeochemical cycles from metabarcoding data

**DOI:** 10.1101/2025.01.30.635649

**Authors:** Arnaud Belcour, Loris Megy, Sylvain Stephant, Caroline Michel, Sétareh Rad, Petra Bombach, Nicole Dopffel, Hidde de Jong, Delphine Ropers

**Author notes:** **To whom correspondence should be addressed. Inria - Université Grenoble Alpes, 655 avenue de l’Europe, Montbonnot, 38334 Saint Ismier CEDEX, France: :** and. **Data availability:** The Tabigecy pipeline is available at https://github.com/ArnaudBelcour/tabigecy. The Python package bigecyhmm and the precomputed EsMeCaTa database are also separately available at https://github.com/ArnaudBelcour/bigecyhmm and https://doi.org/10.5281/zenodo. 13354073, respectively. Datasets used in this article are available at https://doi.org/10.5281/zenodo.14762347. **Competing interests:** The author declare no competing interests.

## Abstract

**Motivation:** Taxonomic analysis of environmental microbial communities is now routinely performed thanks to advances in DNA sequencing. Determining the role of these communities in global biogeochemical cycles requires the identification of their metabolic functions, such as hydrogen oxidation, sulfur reduction, and carbon fixation. These functions can be directly inferred from metagenomics data, but in many environmental applications metabarcoding is still the method of choice. The reconstruction of metabolic functions from metabarcoding data and their integration into coarse-grained representations of geobiochemical cycles remains a difficult bioinformatics problem today.

**Results:** We developed a pipeline, called Tabigecy, which exploits taxonomic affiliations to predict metabolic functions constituting biogeochemical cycles. In a first step, Tabigecy uses the tool EsMeCaTa to predict consensus proteomes from input affiliations. To optimise this process, we generated a precomputed database containing information about 2,404 taxa from UniProt. The consensus proteomes are searched using bigecyhmm, a newly developed Python package relying on Hidden Markov Models to identify key enzymes involved in metabolic function of biogeochemical cycles. The metabolic functions are then projected on coarse-grained representation of the cycles. We applied Tabigecy to two salt cavern datasets and validated its predictions with microbial activity and hydrochemistry measurements performed on the samples. The results highlight the utility of the approach to investigate the impact of microbial communities on geobiochemical processes.

## Introduction

The characterization of environmental microbial communities has enormously progressed thanks to advances in metabarcoding and metagenomics over the past two decades [1]. These approaches are based on high-throughput sequencing and allow the identification of microbial community composition in different environments [2, 3]. Moreover, they enable the analysis of metabolic functions associated with individual microbial species, genera or families in order to understand, for example, their impact on soil biogeochemistry. One of the ultimate objectives is to elucidate the role these organisms play in major biogeochemical cycles [4, 5].

Metagenomics approaches sequence DNA extracted from environmental samples and assemble genomes without the need for lab cultivation [6, 7]. The metabolic functions associated with the genomic data can be directly inferred by matching DNA sequences with corresponding protein sequences in annotated reference databases. In comparison with metagenomics, metabarcoding approaches are directed against specific barcode sequences (*e*.*g*., 16S rRNA in bacteria). The sequences are amplified through Polymerase Chain Reaction (PCR) and compared with a taxonomic reference database to identify the presence of specific taxa in the sample [8]. The association with metabolic functions is then obtained either by literature search or by relying on annotated genomes available in public databases. In metabarcoding, the relation between the DNA extracted from a sample and metabolic functions of the microorganisms is thus more indirect, mediated by taxonomic categories, than in metagenomics. Metabarcoding is still a method of choice in many environmental studies, however, among other things because it (1) requires less DNA for sequencing, (2) is faster when dealing with a great amount of samples, and (3) requires less bioinformatics analysis [9].

The mapping of metabarcoding data to networks of metabolic functions interconverting organic and inorganic compounds present in the environment is a complex bioinformatics problem. Several methods have been developed for extracting metabolic functions from metabarcoding data, such as PICRUSt [10, 11], Paprica [12], Tax4Fun [13, 14], and PanFP [15]. These methods are usually limited to 16S markers and, more fundamentally, they return specific annotation types (EC numbers, KEGG orthologs, …) that may be difficult to interpret and to integrate into aggregated steps of biogeochemical cycles, such as methanogenesis, denitrification, and sulfate reduction. There exist methods for extracting such coarse-grained metabolic functions from sequencing data of microbial communities, notably METABOLIC [16]. This method, however, is restricted to metagenomics data and cannot be used for datasets generated by metabarcoding.

In this work, we propose an integrated approach for the reconstruction of coarse-grained representations of biogeochemical cycles from metabarcoding data which addresses the above problems. The approach takes the form of a bioinformatics pipeline which, in a first step, exploits taxonomic affiliations from metabarcoding sequencing to obtain a list of enzymes (metabolic reactions) occurring in the species associated with the taxa. This step is based on the method EsMeCaTa, which predicts consensus proteomes from taxonomic affiliations [17, 18], and exploits a precomputed database relating taxa to proteomes to speed up computations. In a second step, the list of enzymes (metabolic reactions) inferred from the metabarcoding data are analyzed by means of Hidden Markov Models (HMMs) linking proteins to coarse-grained metabolic functions associated with biogeochemical cycles. This second step is based on METABOLIC [16], while slightly modifying its repertoire of HMMs to account for genes of interest in specific environments. In a third step, the inferred metabolic functions are mapped to carbon, nitrogen, and sulfur cycle diagrams, each consisting of metabolic functions and the compounds they consume and produce. The entire pipeline, called Tabigecy, has been implemented in the workflow language Nextflow [19]. The code of the pipeline as well as the Python package performing the HMM search of metabolic functions (bigecyhmm) and the precomputed EsMeCaTa database are available.

We show the applicability of the Tabigecy pipeline by analyzing two datasets concerning microbial communities in salt caverns [20, 21]. The analysis reproduces the main features of the interpretation of the data in the source publications, but also refines the conclusions and proposes new insights. Our results notably highlight the diversity of metabolic functions active in different caverns, suggest how this diversity could be related to human activity, and discuss the impact of identified metabolic functions on the capacity for CO_2_ and H_2_ storage.

The approach presented in this article is generally applicable to metabarcoding datasets in environmental microbiology, possibly after refining the prediction of coarse-grained metabolic functions to the type of communities and biogeochemical cycles studied. The approach is modular in that the prediction of coarse-grained metabolic functions can also start from a list of proteins obtained from the analysis of metagenomics rather than metabarcoding datasets.

## Results

### Pipeline for inferring coarse-grained representations of biogeochemical cycles from taxonomic affiliations

Tabigecy is a workflow which takes as input taxonomic affiliations resulting from the taxonomic assignment of metabarcoding sequences and which produces as output network diagrams of metabolic functions associated with major biogeochemical cycles (Fig. 1). It accepts the relative abundances of the microorganisms in the community as an additional, optional input to weigh the activity of the metabolic functions. Tabigecy has been implemented in the workflow system Nextflow [19], which allows to construct a computational pipeline from individual tasks defined with input requirements and output declarations. Below we characterize the basic functionality of the pipeline in three steps, putting special emphasis on the novel contributions of this work. For technical details on the implementation of Tabigecy, we refer to the *Methods*.

**Figure 1.**
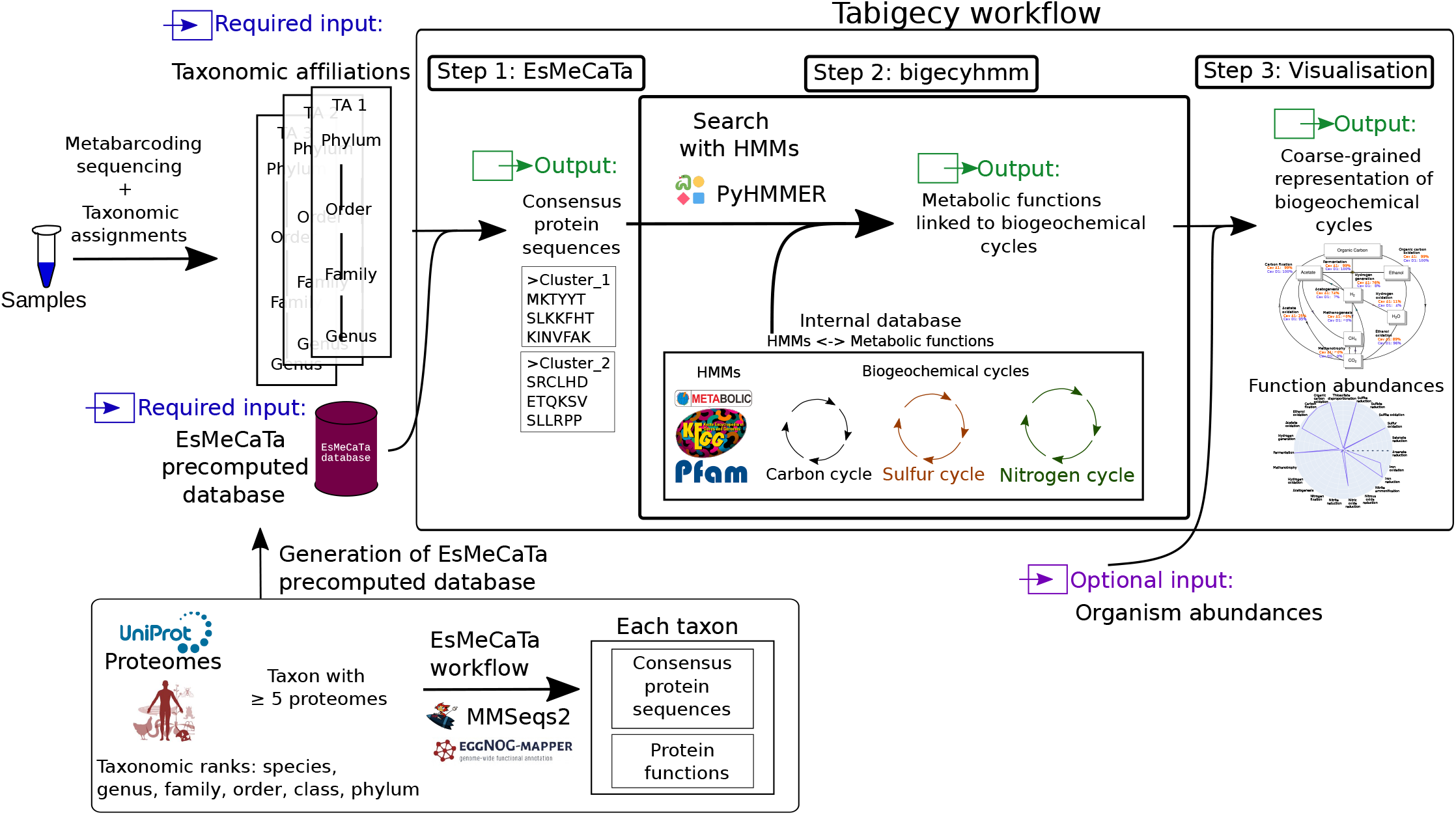
Overview of the Tabigecy pipeline. The input of Tabigecy consists of (1) taxonomic affiliations assigned to metabarcoding data, (2) the EsMeCaTa database with precomputed mappings of concensus proteomes to taxonomic affiliations, and optionally, (3) the abundance of the sequenced organisms in the environmental samples. The output of Tabigecy consists of coarse-grained metabolic functions associated with biogeochemical cycles and their visualisation. The figure also summarizes the generation of the precomputed EsMeCaTa database. The pipeline draws upon a variety of existing bioinformatics tools, and extends and optimizes them for the purpose of this study, as explained in the main text.

#### Prediction of consensus proteomes from taxonomic affiliations

The input of the approach consists of the taxonomic affiliations of the microbial species present in the sample, as identified from the metabarcoding data [22]. We first use EsMeCaTa [17, 18] to predict consensus proteomes from the taxonomic affiliations. In summary, EsMeCaTa searches the UniProt database [23] for the proteomes associated with each taxon (species, genus, family, …). It then uses MMseqs2 [24] to generate consensus sequences for proteins by clustering the UniProt proteomes at a specific taxonomic rank. The resulting consensus proteomes are annotated with eggNOG-mapper [25, 26], which assigns functional categories to the protein clusters based on orthology predictions.

The entire process of EsMeCaTa is resource and time-consuming and may take up to tens of hours to complete. In order to overcome this limitation, we generated a precomputed database of EsMeCaTa predictions and exploit this database to optimize the process. The database was obtained by running EsMeCaTa on 25,874 taxa present in UniProt (Table 1), at different taxonomic ranks, using a computer cluster (*Methods*). This greatly reduces the time required by EsMeCaTa to predict consensus proteomes, typically to tens of minutes for the datasets considered below. Moreover, it improves the reproducibility of the results by ensuring that a well-defined database is used in every run of EsMeCaTa.

**Table 1.** Contents of the precomputed EsMeCaTa database. The column “UniProt taxa” refers to the total number of taxa in UniProt and the column “Included taxa” to those having more than five associated proteomes and included in the precomputed EsMeCaTa database.

| Taxonomic rank | UniProt taxa | Included taxa | Number of proteomes |
| --- | --- | --- | --- |
| Species | 13,187 | 403 | 5,043 |
| Genus | 8,926 | 960 | 11,490 |
| Family | 2,215 | 519 | 9,519 |
| Order | 899 | 292 | 7,169 |
| Class | 373 | 139 | 4,298 |
| Phylum | 274 | 91 | 2,942 |

#### Inference of coarse-grained metabolic functions from consensus proteomes

In a second step, the workflow uses the consensus proteomes derived in the previous step to predict coarse-grained metabolic functions accomplished by environmental microbial communities. We developed the Python package bigecyhmm which implements the search for enzymes involved in metabolic functions using Hidden Markov Models (HMMs), as proposed in METABOLIC [16]. bigecyhmm takes as input protein sequences in FASTA files, which are searched by pyHmmer [27] using an internal database of HMMs. The enzymes thus identified are then mapped to coarsegrained metabolic functions involved in biogeochemical cycles. In comparison with METABOLIC, we slightly extended the repertoire of HMMs to account for metabolic specificities of the microbial communities in salt caverns analysed in this work.

#### Visualisation of metabolic functions in biogeochemical cycles

In a third step, Tabigecy visualises the identified metabolic functions in the form of a polar plot summarizing the major functions. Moreover, it projects the functions on biogeochemical cycle diagrams for carbon, sulfur and nitrogen borrowed from [16]. The diagrams show which organic and inorganic compounds in the environment are consumed and produced by the metabolic functions. If the abundances of the microorganisms in the communities are provided as optional input (*e*.*g*., through high-throughput sequencing of 16S rRNA), then the functions are weighted by the relative abundances. Tabigecy also generates a heatmap showing, for each metabolic function associated with an HMM, the relative abundance of the microorganisms (when available) or the relative occurrence of this function in the taxa identified in the sample (*Methods*).

### Application of the pipeline to the analysis of salt cavern communities

Salt caverns are artificial underground formations that are created by drilling a well into a salt formation and injecting water to dissolve the salt, called solution mining. The resulting brine, consisting of water mixed with salt, is extracted, leaving space for storage. The impermeability of salt caverns prevents gas leakage, making them ideal for gas storage [28]. Despite the high salinity of the environment, microbial life, mostly bacteria and archaea, is abundant in salt caverns. In case of hydrogen storage, microbial activity can change the gas composition and damage the operating infrastructure [29]. Current knowledge of the microbiology of salt caverns is limited for a number of reasons, including the difficulty to obtain sufficient DNA from samples for characterising microbial diversity, the under-representation of salt cavern communities in public genome databases, and the difficulty of culturing microbial species present in caverns in the laboratory [30, 31]. Predicting the metabolic functions performed by microbial communities in salt caverns is helpful for designing mitigation strategies against harmful microbial reactions. We therefore applied our Tabigecy pipeline to two publicly available datasets (*Methods*).

#### Diversity of acetate metabolism in salt caverns

[21] investigated the microbial diversity in five German salt caverns (Cav. A to E) which have been used for natural gas storage. Taxonomic assignment of the metabarcoding reads was performed on duplicate samples from the caverns and the resulting affiliations were used to predict metabolic functions using Tabigecy (Fig. 1). The results for each replicate and each sample are shown in Supplementary Figs S1 and S7. As observed by [21] at the genomic level, the replicates of Cav. A, C, and E yield reproducible functional profiles, whereas replicates of Cav. B and D have several differences.

We performed a Principal Component Analysis (PCA) on the predicted metabolic functions in the different samples and found that the caverns separate into two groups (Supplementary Fig. S2A). We analyzed one representative sample of each group in detail (A1 and D1, the first replicates of caverns A and D). The polar plot in Fig. 2A shows the main metabolic functions performed by the communities in the two caverns. The plot reveals that many functions are present in both caverns, for example generic metabolic functions like fermentation, organic carbon oxidation, and carbon fixation, but also more specific functions such as sulfite reduction and sulfur oxidation. However, the analysis also predicts differences, such as the presence of hydrogen generation in A1, but not in D1. One of the major differences concerns acetate metabolism: whereas acetate is produced in A1, it is consumed in D1. The latter observation is confirmed by the PCA in Supplementary Fig. S2, where the analysis of the main principal components shows that acetate oxidation and acetogenesis are seen to be strongly but negatively correlated.

**Figure 2.**
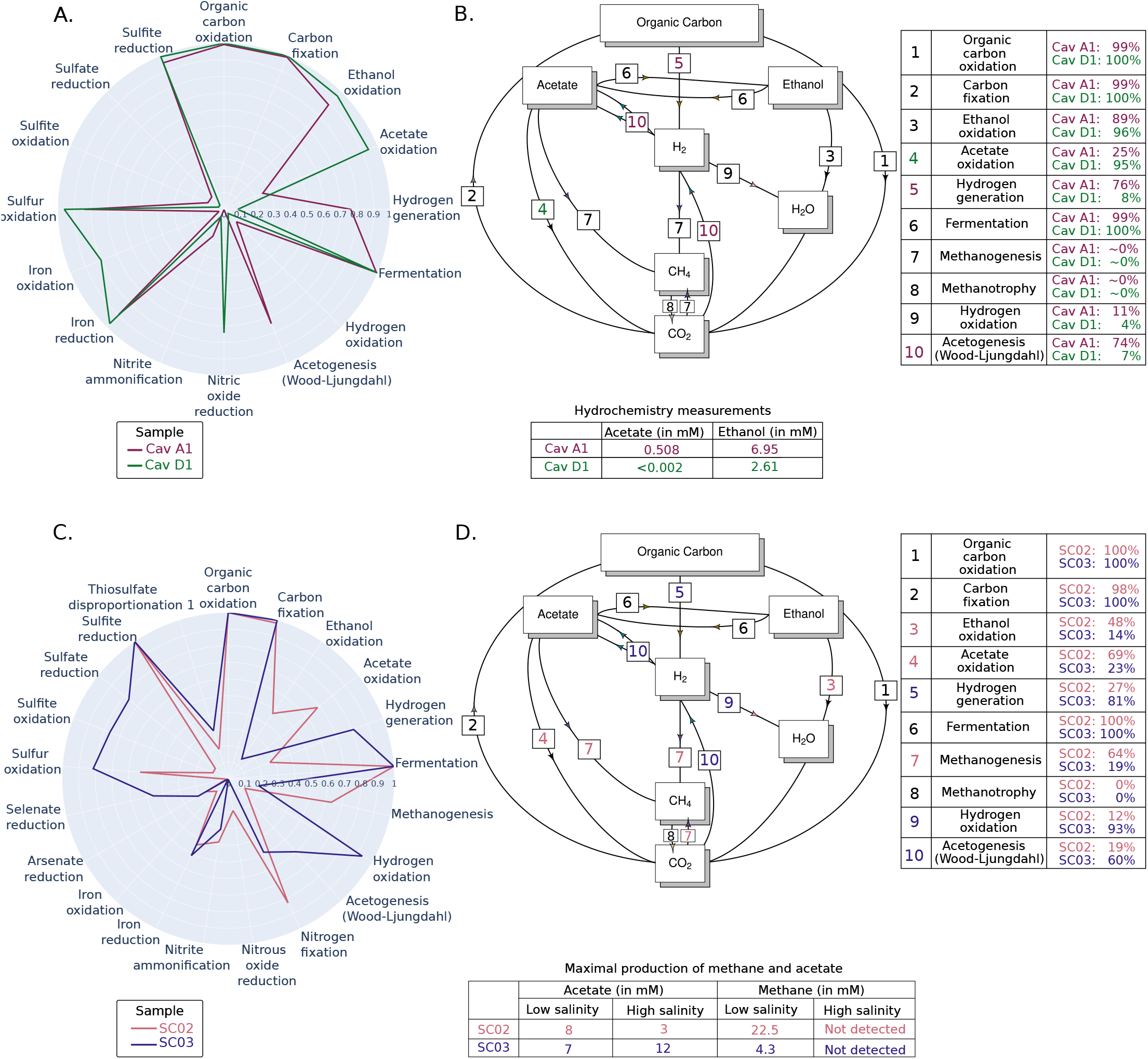
Reconstruction of coarse-grained representations of biogeochemical cycles from the taxonomic affiliations of salt cavern communities. Salt cavern datasets from [21] and [20] were analysed using the Tabigecy pipeline. **A**. Polar plot indicating for each major metabolic function the relative abundance of microorganisms found in two samples (A1 and D1) in the Schwab dataset. Only functions having at least 10% of relative abundance in a sample are shown. **B**. Projection of the metabolic functions from panel A on the carbon cycle diagram from [16], with weights in the functions given by the relative abundance of the organisms in the two samples. Measurements of acetate and ethanol from [21] are shown in the accompanying table. **C**. As in panel A, but for two samples (SC02 and SC03) from the Bordenave dataset. **D**. As in panel B, but for the Bordenave dataset, completed with measurements of acetate and methane in high and low salinity conditions [20].

The predicted metabolic functions were mapped to the biogeochemical cycle diagram for carbon, shown in Fig. 2B. The functions were weighted with the measured relative abundances of microorganisms predicted to perform these functions, based on the information provided by the consensus proteomes. When focusing on acetate production and consumption, the diagram indicates that acetogenesis from hydrogen and CO_2_, *via* the Wood-Ljungdahl pathway, occurs in A1, whereas acetate oxidation takes place in D1. These two predictions are strongly supported by the abundance data, which suggest that the vast majority of microorganisms in A1 have the capacity for acetogenesis (74%) and in D1 for acetate oxidation (95%). In both caverns, some ethanol oxidation, that is, the conversion of ethanol and CO_2_ to acetate, is predicted to occur.

The above predictions can be validated by hydrochemical measurements reported in the same study [21]. Acetate is present at a concentration of 0.5 mM in A1, but is undetectable in D1, consistent with the predominance of acetate production in A1 and of acetate consumption in D1. Ethanol is present in both samples, at high concentrations (2.6 and 7.0 mM). Ethanol is probably artificially introduced in caverns due to injection of workover fluids, showing the possible consequences of human intervention on biogeochemical cycles.

The reconstruction of coarse-grained representations of biogeochemical cycles by means of Tabigecy is thus shown to reproduce and refine the main observations in the original study. In particular, our analysis provides further evidence for the hypothesis of Schwab *et al*. that the community is capable of producing acetate from hydrogen and CO_2_. While Fig. 2 focuses on the carbon cycle, Supplementary Fig. S3 displays the projection of the metabolic functions on the nitrogen and sulfur cycles.

#### Impact of microbial activity on CO_2_ storage capacity

[20] analysed four samples from a salt cavern plant in Canada, including brine samples obtained from two salt caverns (SC02 and SC03), in view of the use of these caverns for CO_2_ storage. The two caverns differ in that one had been used for oily sand disposal (SC02) whereas the other was unused (SC03). The authors already assigned taxonomic categories to the data and provided the relative abundances of the microorganisms in the samples, which were used as input for Tabigecy. The results for all samples are shown in Supplementary Figs S4 and S8.

Like for the previous salt cavern study, we find a global correspondence of metabolic functions in the two samples but also interesting differences (Fig. 2C). A first observation is that in both samples, as observed by [21], a vast majority of organisms are capable of fermentation, organic carbon oxidation, carbon fixation, and sulfite reduction. Moreover, the samples show predicted differences in acetate metabolism, like in the previous study. Whereas acetate oxidation (consumption) is found in SC02, acetate is produced in the cavern corresponding to SC03. Other major differences between the two samples are the capacity for methanogenesis, present in SC02 (64% of microorganisms) but much less so in SC03 (19%), and the capacity for hydrogen oxidation, present in SC03 (93%) but much less so in SC02 (12%). This observation is confirmed by the differential correlation of the metabolic functions with the first dimension of the PCA, which accounts for almost 80% of the variance in the data (Supplementary Fig. S5).

Mapping the identified functions to the carbon cycle diagram suggests possible explanations for the above-mentioned differences (Fig. 2D). It seems that in SC02, hydrogen accumulating in the caverns is mostly converted into methane, whereas in SC03, hydrogen is utilized both by acetogenesis and hydrogen oxidation. This hypothesis is partially validated by the microbial activity measurements carried out through enrichment cultures in the laboratory, in the presence of CO_2_ and for different levels of salinity [20]. A four-fold higher concentration of methane was found in SC02 as compared to SC03 (at low salinity), while acetate is detected at higher concentrations in SC03 as compared to SC02 (at high salinity). These differences are expected given the higher capacity for methanogenesis in SC02 and for acetogenesis in SC03. Given the high salinity in salt caverns, the absence of methanogenesis in high-salinity conditions suggests little consumption of CO_2_, a substrate for methanogenesis together with hydrogen. This implies a lower risk for CO_2_ loss in case the caverns are used for CO_2_ storage.

The above observations concern the carbon cycle, but analysis of the nitrogen and sulfur cycles shown in Supplementary Fig. S6 may provide further insights. The predicted strong presence of sulfite reduction, producing toxic, corrosive H_2_S gas, and other sulfur-related functions is an interesting observation for further research on the microbiology of salt caverns [29].

## Discussion

Tabigecy is a workflow to predict coarse-grained metabolic functions accomplished by microbial communities from metabar- coding data and map these functions to global biogeochemical cycles. Tabigecy takes as input taxonomic affiliations inferred from the sequence data and predicts consensus proteomes at the most specific taxonomic ranks possible given the available proteomes in public databases. The consensus proteomes are then analyzed to identify metabolic functions associated with key enzymes and these functions are projected on global cycles for carbon, nitrogen, and sulfur. The identified metabolic functions can be given more or less weight by taking into account, as optional input, the measured abundances of the microorganisms predicted to carry out the functions.

The pipeline is built upon existing bioinformatics tools but adapts and optimises them for our purpose. Tabigecy uses EsMeCaTa for the prediction of consensus proteomes from taxonomic affiliations [17, 18], but exploits a newly developed precomputed database which strongly reduces run times (around 60-fold for the datasets considered here, with half the number of the cores, see *Methods*). While the idea of inferring metabolic functions from sequence data and the projection of these functions on diagrams of biogeochemical cycles was borrowed from METABOLIC [16], Tabigecy adapts the approach both on the conceptual and implementational level. Instead of taking metagenomics data as input, we start from metabarcoding data. The latter are widely used in studies of environmental communities where the DNA yields of samples are often low. Moreover, we reimplemented one of the core tasks of METABOLIC, the HMM search of proteomes to identify enzymes involved in metabolic functions, in a stand-alone Python package, bigecyhmm, allowing this task to be integrated into Tabigecy and other pipelines. We also slightly enlarged the repertoire of HMMs for the purpose of the specific communities analysed in this work.

We tested the capabilities of Tabigecy by analyzing two biogeochemistry datasets obtained in salt caverns. These caverns can be used for the storage of hydrogen and CO_2_, which poses the question of the interactions between the stored gases and resident microbial communities. The application of Tabigecy reproduced the observations made in the original source publications [20, 21], but also provided a more comprehensive and quantitative view. Instead of limiting the analysis to a few *a-priori* selected metabolic functions, we systematically scanned the datasets for evidence of the presence of dozens of functions. Moreover, the integration of measurements of microbial abundances makes it possible to assign a quantitative weight to predicted metabolic functions. It is important to emphasize that the predicted functions correspond to a metabolic potential but not necessarily to a metabolic activity, as the enzymes involved in the functions may be inactive. Some of the predictions of Tabigecy, however, could be validated by relating the microbial functions to reported measurements of substrates and products obtained *in situ* or from laboratory enrichment cultures.

Tabigecy is directly applicable to the study of other environmental microbial communities, for which it is often difficult to extract enough DNA for metagenomic analyses. This generalization of the pipeline to other applications may require the extension of the repertoire of available HMMs and metabolic functions. Moreover, one needs to be aware of possible limitations and biases that were also encountered in the analysis of the salt cavern datasets. First, the functional characterization of communities with little-studied microorganisms may be less reliable. When few relatively close proteomes are available, EsMeCaTa will use more distant organisms to produce consensus proteomes [18]. The latter are potentially less representative for the microorganisms under study. In addition, due to evolutionary divergence, bigecyhmm may have more difficulty in identifying, in the proteomes, key enzymes of metabolic functions. A second bias comes with the use of 16S barcode sequences to quantify abundances of specific microorganisms in a sample [32]. The copy number of 16S genes varies within and between species [33], which may lead to over- or underestimated abundances. While methods have been developed to correct for this bias by predicting 16S gene copy numbers [11], they may be difficult to apply in the case of poorly studied microorganisms.

A major strength of Tabigecy is that it integrates the metabolic functions inferred from the sequence data into diagrams that provide a coarse-grained representation of biogeochemical cycles. These diagrams are helpful for understanding the functioning of the communities and identifying possible perturbations due to human interference [34], as illustrated by the analysis of the Schwab dataset. The use of the diagrams could be pushed further by realizing that every identified coarsegrained metabolic function corresponds to a macroreaction, *e*.*g*., 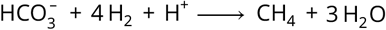 for methanogenesis. Together, the reactions describe the flow of material through carbon, nitrogen, and sulfur cycles. The network of metabolic macroreactions thus obtained could be transformed into a mathematical model enabling the quantitative analysis and simulation of the community dynamics [35]. This opens the perspective of gaining a better, time-resolved understanding of the interactions between human activities and environmental microbial communities, for example in the context of underground gas storage and other industrial applications.

## Methods

### Implementation of the Tabigecy pipeline

The Tabigecy pipeline was developed as a Nextflow workflow [19]. The pipeline can be directly called at its GitHub repository using *next*fl*ow run ArnaudBelcour/tabigecy* … in the local command line, as detailed in the README file of Tabigecy. Tabigecy consists of three processes (Fig. 1) and expects three mandatory inputs: (1) a file with taxonomic affiliations in tabulated format, processed by the ETE3 package [36], (2) the precomputed EsMeCaTa database, and (3) an output folder. There are two optional arguments: (1) a tabulated file listing the abundances associated with the taxonomic affiliations and (2) the number of cores to be used by bigecyhmm.

### Generation and use of the EsMeCaTa database

To obtain the input for the construction of the EsMeCaTa database, a SPARQL query was performed on the UniProt Endpoint [23], resulting in a list of all taxa available for each taxonomic rank (‘species’, ‘genus’, ‘family’, ‘order’, ‘class’ and ‘phylum’), corresponding to six tabulated files. For each file, the ‘esmecata proteomes’ command was run against UniProt (version 2024_05; [23]), using the rank limit option associated with the corresponding taxonomic rank. For example, when processing species, the rank limit parameter of EsMeCaTa was set to ‘species’, to avoid EsMeCaTa using higher taxonomic ranks. EsMeCaTa produces a list of taxa, having at least five associated proteomes, and the corresponding proteomes. The latter were then clustered with ‘esmecata clustering’ using 36 CPUs on a HPC cluster by means of MMseqs2 (version 15.6f452; [24]). Next, the consensus proteomes predicted by EsMeCaTa were annotated by means of ‘esmecata annotation’ using eggNOG-mapper (version 2.1.12) and the eggNOG database (version 5.0.2; [25, 26]). The generation of the annotated consensus proteomes using the HPC cluster took 104 hours for ‘esmecata proteomes’, 25 hours for ‘esmecata clustering’ (32 cores), and 31 hours for ‘esmecata annotation’ (32 cores).

The folders containing the above results were then used as input by the new EsMeCaTa command *esmecata_create_db*. The command compiles the information for each taxon into a consensus proteome FASTA file and an annotation file. The annotation file is generated by combining the results from eggNOG-mapper and MMseqs2, listing for each protein cluster the annotation and the included proteins. The new subcommand *esmecata precomputed* extends the standard *esmecata work*fl*ow* command to exploit the database thus obtained. It takes as input a tabulated file containing taxonomic affiliations and the compressed zip archive of the database. EsMeCaTa parses the taxonomic affiliations using ETE3 and searches for the presence of corresponding taxa in the database. If it finds a match, the consensus proteome FASTA file and the annotation file are extracted and transferred to the output folder.

The above commands and the precomputed database were included in a new version of EsMeCaTa (version 0.6.0).

### Implementation of the bigecyhmm package

We developed the Python package bigecyhmm, which re-implements the task in METABOLIC [16] concerned with the prediction of coarse-grained metabolic functions in biogeochemical cycles from consensus protein sequences. The implementation accompanying the original METABOLIC approach is a complex pipeline starting from the reads in metagenomics datasets. The metabolic function prediction task has multiple dependencies that do not allow this task to be easily isolated from the pipeline. The bigecyhmm package was designed to be a portable, self-contained, and easy-to-use package.

Function prediction is called by means of the command *bigecyhmm* and takes as input either (1) a FASTA file with protein sequences or (2) a folder containing multiple such FASTA files. An internal database, contained within the package, includes the HMM profiles for enzymes associated with biogeochemical cycles used by METABOLIC. Compared to the profiles in METABOLIC, several HMM profiles associated with the following genes were added: *hycE* (formate lyase), *acsB* (acetyl-CoA synthase), *fthfs/fhs* (formate–tetrahydrofolate ligase), *mtaB/mtbA* (coenzyme M methyltransferase), and *hydB/hydG* (sulfhydrogenase).

The input protein sequences are searched for matches with the HMM profiles using PyHMMER [27], a Python library binding to HMMER [37]. Boolean functions consisting of combinations of OR and AND operators between HMM profiles are used to conclude on the presence of specific coarsegrained metabolic functions. When a folder of FASTA files is provided as input to bigecyhmm, the number of cores can be specified as an optional argument of the command. bigecyhmm then parallelises the process by distributing the HMM search of individual protein sequence files over the cores.

The biogeochemical cycle diagrams of [16] were reconstructed as svg templates and the metabolic function predictions are projected on the diagrams using the Pillow library [38]. bigecyhmm also computes the occurrence and, when a file with the abundances of microbial species is given as an additional input, the relative abundance of each metabolic function. Occurrences are defined as the number of taxa possessing a function as a fraction of the total number of taxa in the community. Relative abundances correspond to the sum of the abundances of all taxa possessing a function divided by the sum of abundances in the sample. Heatmaps describing either the occurrence or abundance of all metabolic functions are generated using seaborn [39] and matplotlib [40, 41] packages. The occurrences and abundances of major identified functions in the biogeochemical cycles are also shown in polar plots generated by means of the plotly package [42].

### Salt cavern datasets

The communities sequenced in the study of [21] were analysed by first retrieving the reads available in European Nucleotide Archive (ENA, accession number PRJEB49822). The more than 500,000 reads were processed by means of the FROGs pipeline (version 4.1.0; [43, 44]) accessible through Galaxy France (https://usegalaxy.fr/; [45]). In more detail, reads were first merged with VSEARCH (version 2.17.0; [46]) and dereplicated. They were then clustered using swarm (version 3.0.0; [47]). Chimera were identified by means of VSEARCH and removed. Only clusters comprising at least 0.005% of all sequences were retained and those aligning to contaminants, as determined by means of BLAST (version 2.10; [48]) were removed. Taxonomic assignment was then performed using BLAST and the reference database 16S SILVA 138.1 [49]. The resulting file in Biological Observation Matrix (BIOM) format was converted into a tabulated file.

Table 2 in the study of [20] was used to create the Tabigecy input files with taxonomic affiliations. Each genus in the table was processed by the ETE3 package (version 3.1.3; [36]) to retrieve the whole lineage (from genus to kingdom). Associating a genus with its complete lineage is necessary to prepare the input file for EsMeCaTa, in order to allow EsMeCaTa to use a higher taxonomic level when not enough proteomes are found at the genus level.

### Application of the pipeline to salt cavern communities

The two salt cavern datasets were processed using Tabigecy (version 0.1.1). Consensus proteome predictions were performed by EsMeCaTa (version 0.6.0) with the precomputed database (version 1.0.0) generated from UniProt release 2024_05 and the NCBI Taxonomy database with timestamp 2024-10-01. Prediction and visualisation of the coarse-grained metabolic functions was achieved by means of bigecyhmm (version 0.1.5). On a personal computer, using 5 cores, the whole workflow took 14 minutes to complete for the Schwab dataset and 3 minutes for the Bordenave dataset. By comparison, the original EsMeCaTa application without the precomputed database took 12.5 hours to process the Schwab dataset and 3 hours for the Bordenave dataset on a computer cluster with 10 cores. The PCA biplots in the Supplementary information were generated using R (version 4.4.1; [50]), factoextra library (version 1.0.7; [51]) and the ade4 library (version 1.7-22; [52, 53, 54, 55]). Correlation plots for PCA dimensions and function abundances were created by means of the R corrplot library (version 0.94; [56]).

## Acknowledgments

Most computations for this work were performed using the GRICAD infrastructure in Grenoble (https://gricad.univ-grenoblealpes.fr).

## Author contributions statement

A.B. conceived the method and implemented the pipeline. A.B., L.M. and D.R. generated the pre-computed EsMeCaTa database. A.B., H.d.J. and D.R. conceived the experiments. A.B. conducted the experiments. A.B., S.S., C.M., S.R., P.B., N.D., H.d.J. and D.R. analysed the results. A.B., H.d.J. and D.R. wrote the manuscript and all authors reviewed the manuscript.

## References

[1] Yishay Pinto and Ami S. Bhatt. “Sequencing-based analysis of microbiomes”. In: Nat Rev Genet 25.12 (2024), pp. 829–45.

[2] Luke R. Thompson, Jon G. Sanders, Daniel McDonald, et al. “A communal catalogue reveals Earth’s multiscale microbial diversity”. In: Nature 551.7681 (2017), pp. 457–63.

[3] Justin P. Shaffer, Louis-Félix Nothias, Luke R. Thompson, et al. “Standardized multi-omics of Earth’s microbiomes reveals microbial and metabolite diversity”. In: Nat Microbiol 7.12 (2022), pp. 2128–50.

[4] Shinichi Sunagawa, Silvia G. Acinas, Peer Bork, et al. “Tara Oceans: towards global ocean ecosystems biology”. In: Nat Rev Microbiol 18.8 (2020), pp. 428–45.

[5] Noah W. Sokol, Eric Slessarev, Gianna L. Marschmann, et al. “Life and death in the soil microbiome: how ecological processes influence biogeochemistry”. In: Nat Rev Microbiol 20.7 (2022), pp. 415–30.

[6] Kevin Chen and Lior Pachter. “Bioinformatics for whole-genome shotgun sequencing of microbial communities”. In: PLoS Comput Biol 1.2 (2005), e24.

[7] Alejandra Escobar-Zepeda, Arturo Vera-Ponce de León, and Alejandro Sanchez-Flores. “The road to metagenomics: from microbiology to DNA sequencing technologies and bioinformatics”. In: Front Genet 6 (2015), p. 348.

[8] Pierre Taberlet, Eric Coissac, François Pompanon, et al. “Towards next-generation biodiversity assessment using DNA metabarcoding”. In: Mol Ecol 21.8 (2012), pp. 2045–50.

[9] Meghana Srinivas, Orla O’Sullivan, Paul D. Cotter, et al. “The application of metagenomics to study microbial communities and develop desirable traits in fermented foods”. In: Foods 11.20 (2022), p. 3297.

[10] Morgan G. I. Langille, Jesse Zaneveld, J. Gregory Caporaso, et al. “Predictive functional profiling of microbial communities using 16S rRNA marker gene sequences”. In: Nat Biotechnol 31.9 (2013), pp. 814–21.

[11] Gavin M. Douglas, Vincent J. Maffei, Jesse R. Zaneveld, et al. “PICRUSt2 for prediction of metagenome functions”. In: Nat Biotechnol 38.6 (2020), pp. 685–8.

[12] Jeff S. Bowman and Hugh W. Ducklow. “Microbial communities can be described by metabolic structure: a general framework and application to a seasonally variable, depth-stratified microbial community from the coastal West Antarctic peninsula”. In: PLoS One 10.8 (2015), e0135868.

[13] Kathrin P. Aßhauer, Bernd Wemheuer, Rolf Daniel, et al. “Tax4Fun: predicting functional profiles from metagenomic 16S rRNA data”. In: Bioinformatics 31.17 (2015), pp. 2882–4.

[14] Franziska Wemheuer, Jessica A. Taylor, Rolf Daniel, et al. “Tax4Fun2: prediction of habitat-specific functional profiles and functional redundancy based on 16S rRNA gene sequences”. In: Environ Microbiome 15.1 (2020), p. 11.

[15] Se-Ran Jun, Michael S. Robeson, Loren J. Hauser, et al. “PanFP: pangenome-based functional profiles for microbial communities”. In: MC Res Notes 8.1 (2015), p. 479.

[16] Zhichao Zhou, Patricia Q. Tran, Adam M. Breister, et al. “METABOLIC: high-throughput profiling of microbial genomes for functional traits, metabolism, biogeochemistry, and community-scale functional networks”. In: Microbiome 10.1 (2022), p. 33.

[17] Baptiste Ruiz, Arnaud Belcour, Samuel Blanquart, et al. “SPARTA: Interpretable functional classification of microbiomes and detection of hidden cumulative effects”. In: PLoS Comput Biol 20.11 (2024), e1012577.

[18] Arnaud Belcour, Pauline Hamon-Giraud, Alice Mataigne, et al. “Estimating consensus proteomes and metabolic functions from taxonomic affiliations”. In: bioRxiv (2025).

[19] Paolo Di Tommaso, Maria Chatzou, Evan W. Floden, et al. “Nextflow enables reproducible computational workflows”. In: Nat Biotechnol 35.4 (2017), pp. 316–9.

[20] Sylvain Bordenave, Indranil Chatterjee, and Gerrit Voordouw. “Microbial community structure and microbial activities related to CO2 storage capacities of a salt cavern”. In: Int Biodeterior Biodegradation 81 (2013), pp. 82–7.

[21] Laura Schwab, Denny Popp, Guido Nowack, et al. “Structural analysis of microbiomes from salt caverns used for underground gas storage”. In: Int J Hydrogen Energy 47.47 (2022), pp. 20684–94.

[22] Yong-Xin Liu, Yuan Qin, Tong Chen, et al. “A practical guide to amplicon and metagenomic analysis of microbiome data”. In: Protein & Cell 12.5 (2021), pp. 315–30.

[23] The UniProt Consortium. “UniProt: the universal protein knowledgebase in 2025”. In: Nucleic Acids Res 53.D1 (2025), pp. D609–17.

[24] Martin Steinegger and Johannes Söding. “MMseqs2 enables sensitive protein sequence searching for the analysis of massive data sets”. In: Nat Biotechnol 35.11 (2017), pp. 1026–8.

[25] Jaime Huerta-Cepas, Damian Szklarczyk, Davide Heller, et al. “eggNOG 5.0: a hierarchical, functionally and phylogenetically annotated orthology resource based on 5090 organisms and 2502 viruses”. In: Nucleic Acids Res. 47.D1 (2019), pp. D309–14.

[26] Carlos P Cantalapiedra, Ana Hernández-Plaza, Ivica Letunic, et al. “eggNOG-mapper v2: Functional annotation, orthology assignments, and domain prediction at the metagenomic scale”. In: Mol Biol Evol 38.12 (2021), pp. 5825–9.

[27] Martin Larralde and Georg Zeller. “PyHMMER: a Python library binding to HMMER for efficient sequence analysis”. In: Bioinformatics 39.5 (2023), btad214.

[28] Danae A. Voormeij and George J. Simandl. “Geological, ocean, and mineral CO2 sequestration options: a technical review”. In: Geosci Can (2004).

[29] Nicole Dopffel, Stefan Jansen, and Jan Gerritse. “Microbial side effects of underground hydrogen storage – Knowledge gaps, risks and opportunities for successful implementation”. In: Int J Hydrogen Energy 46.12 (2021), pp. 8594–606.

[30] A. López-López, M. Richter, A. Peña, J. Tamames, et al. “New insights into the archaeal diversity of a hypersaline microbial mat obtained by a metagenomic approach”. In: Syst Appl Microbiol 36.3 (2013), pp. 205–14.

[31] Mirosław Słowakiewicz, Weronika Goraj, Tomasz Segit, et al. “Hydrochemical gradients driving extremophile distribution in saline and brine groundwater of southern Poland”. In: Environ Microbiol Rep 16.5 (2024), e70030.

[32] Monica Steffi Matchado, Malte Rühlemann, Sandra Reitmeier, et al. “On the limits of 16S rRNA gene-based metagenome prediction and functional profiling”. In: Microb. Genom. 10.2 (2024), p. 001203.

[33] Silvia G. Acinas, Luisa A. Marcelino, Vanja Klepac-Ceraj, et al. “Divergence and redundancy of 16S rRNA sequences in genomes with multiple rrn operons”. In: J Bacteriol 186.9 (2004), pp. 2629–35.

[34] K.K. Amundson, M.A. Borton, and M.J. Wilkins. “Anthropogenic impacts on the terrestrial sub-surface biosphere”. In: Nat Rev Microbiol (2024).

[35] S. Widder, R.J. Allen, T. Pfeiffer, et al. “Challenges in microbial ecology: building predictive understanding of community function and dynamics”. In: ISME J. 10.11 (2016), pp. 2557–68.

[36] Jaime Huerta-Cepas, François Serra, and Peer Bork. “ETE 3: Reconstruction, analysis, and visualization of phylogenomic data”. In: Mol Biol Evol 33.6 (2016), pp. 1635–8.

[37] Sean R. Eddy. “Accelerated profile HMM searches”. In: PLoS Comput Biol 7.10 (2011), e1002195.

[38] Andrew Murray, Hugo van Kemenade wiredfool, et al. python-pillow/Pillow: 11.1.0. Version 11.1.0. 2025.

[39] Michael L. Waskom. “seaborn: statistical data visualization”. In: J Open Source Softw 6.60 (2021), p. 3021.

[40] J. D. Hunter. “Matplotlib: A 2D graphics environment”. In: Comput. Sci. Eng. 9.3 (2007), pp. 90–5.

[41] The Matplotlib Development Team. Matplotlib: Visualization with Python. Version v3.10.0. 2024.

[42] Plotly Technologies Inc. Collaborative data science. Montreal, QC, 2015. URL: https://plot.ly.

[43] Frédéric Escudié, Lucas Auer, Maria Bernard, et al. “FROGS: Find, rapidly, OTUs with Galaxy solution”. In: Bioinformatics 34.8 (2018), pp. 1287–94.

[44] Maria Bernard, Olivier Rué, Mahendra Mariadassou, et al. “FROGS: a powerful tool to analyse the diversity of fungi with special management of internal transcribed spacers”. In: Brief Bioinform 22.6 (2021), bbab318.

[45] The Galaxy Community. “The Galaxy platform for accessible, reproducible, and collaborative data analyses: 2024 update”. In: Nucleic Acids Res 52.W1 (2024), W83–94.

[46] Torbjørn Rognes, Tomáš Flouri, Ben Nichols, et al. “VSEARCH: a versatile open source tool for metagenomics”. In: PeerJ 4 (2016), e2584.

[47] Frédéric Mahé, Lucas Czech, Alexandros Stamatakis, et al. “Swarm v3: towards tera-scale amplicon clustering”. In: Bioinformatics 38.1 (2021), pp. 267–9.

[48] Christiam Camacho, George Coulouris, Vahram Avagyan, et al. “BLAST+: architecture and applications”. In: BMC Bioinform 10 (2009), p. 421.

[49] Christian Quast, Elmar Pruesse, Pelin Yilmaz, et al. “The SILVA ribosomal RNA gene database project: improved data processing and web-based tools”. In: Nucleic Acids Res 41 (2013), pp. D590– 6.

[50] R Core Team. R: A Language and Environment for Statistical Computing. R Foundation for Statistical Computing. Vienna, Austria, 2024. URL: https://www.R-project.org/.

[51] Alboukadel Kassambara and Fabian Mundt. factoextra: Extract and Visualize the Results of Multivariate Data Analyses. R package version 1.0.7. 2020.

[52] Daniel Chessel, Anne-Béatrice Dufour, and Jean Thioulouse. “The ade4 Package – I: one-table methods”. In: R News 4.1 (2004), pp. 5–10.

[53] Stéphane Dray, Anne-Béatrice Dufour, and Daniel Chessel. “The ade4 Package – II: two-table and K-table methods”. In: R News 7.2 (2007), pp. 47–52.

[54] Stéphane Dray and Anne-Béatrice Dufour. “The ade4 package: implementing the duality diagram for ecologists”. In: J Stat Softw 22 (2007), pp. 1–20.

[55] Jean Thioulouse, Stéphane Dray, Anne-Béatrice Dufour, et al. Multivariate Analysis of Ecological Data with ade4. New York, NY: Springer, 2018.

[56] Taiyun Wei and Viliam Simko. R package ‘corrplot’: Visualization of a correlation matrix. (Version 0.94). 2024.

