## Supplementary File 1 for "Predicting coarse-grained representations of biogeochemical cycles from metabarcoding data"

### Supplementary information for "Predicting coarse-grained representations of biogeochemical cycles from metabarcoding data"

**Arnaud Belcour**[1,2✉], **Loris Megy**[3], **Sylvain Stephant**[4], **Caroline Michel**[4], **Sétareh Rad**[4], **Petra Bombach**[5], **Nicole Dopffel**[6], **Hidde de Jong**[1,2], **Delphine Ropers**[1,2✉]

[1]Univ. Grenoble Alpes, Inria, 38000 Grenoble, France; [2]Université Grenoble Alpes, CNRS, LIPhy, Grenoble, France; [3]Gricad, Inria, CNRS, Université Grenoble Alpes, Grenoble INP, 38000 Grenoble, France; [4]French Geological Survey (BRGM), Orléans, France; [5]Isodetect GmbH, Germany; [6]NORCE Norwegian Research Center AS, Norway

✉ To whom correspondence should be addressed. Inria - Université Grenoble Alpes, 655 avenue de l'Europe, Montbonnot, 38334 Saint Ismier CEDEX, France:

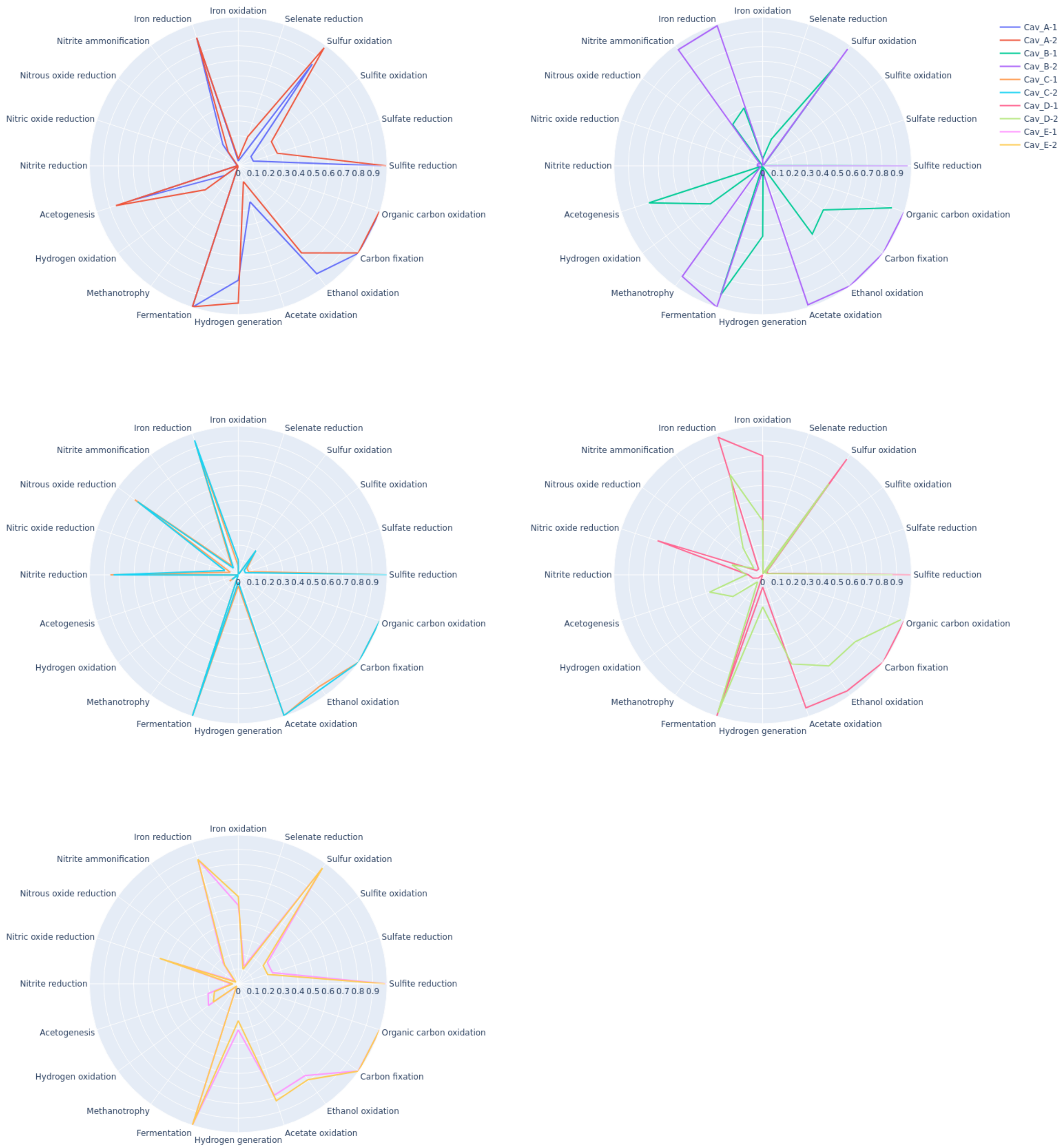


**Figure 1.** Polar plots showing, for each major metabolic function, the relative abundance of microorganisms found in the samples in the Schwab dataset [1]. Only functions having at least 10% of relative abundance in at least one of the samples are shown.

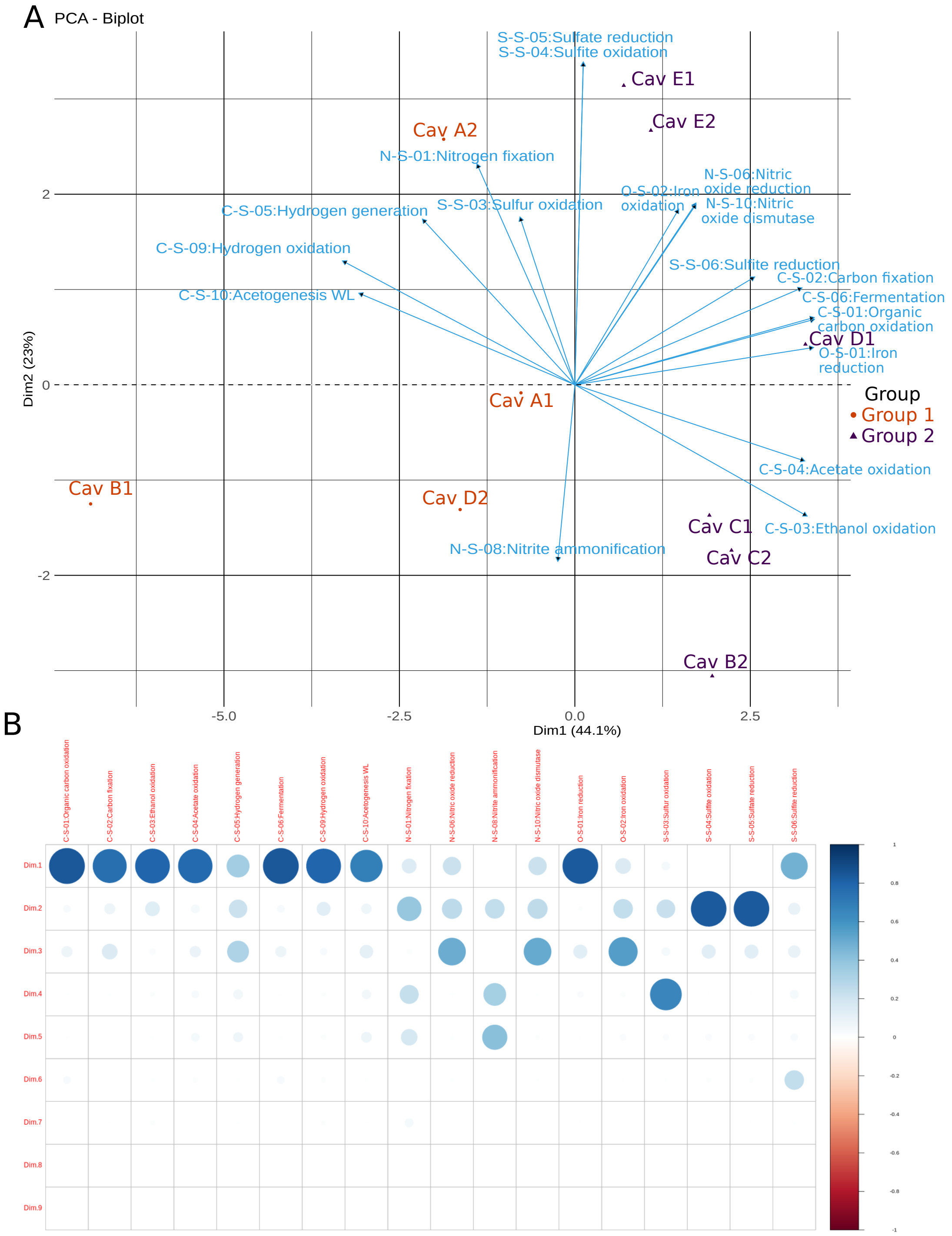


**Figure 2.** Principal Component Analysis (PCA) of the results obtained for the Schwab dataset. A. Projection on the first two PCA dimensions of the relative abundances of metabolic functions reconstructed from the samples in the Schwab dataset [1]. The vectors in the biplot represent the metabolic functions and the points the salt caverns. The latter are clustered in two groups located on opposite sides of the origin: one group for A1, A2, B1 and D2, and one group for the remaining caverns. Negative correlations are observed between metabolic functions. For example, the vectors representing acetogenesis and acetate oxidation point in opposite directions. B. Plot of the correlation of the relative abundances of metabolic functions in the Schwab dataset samples with the 9 PCA dimensions.

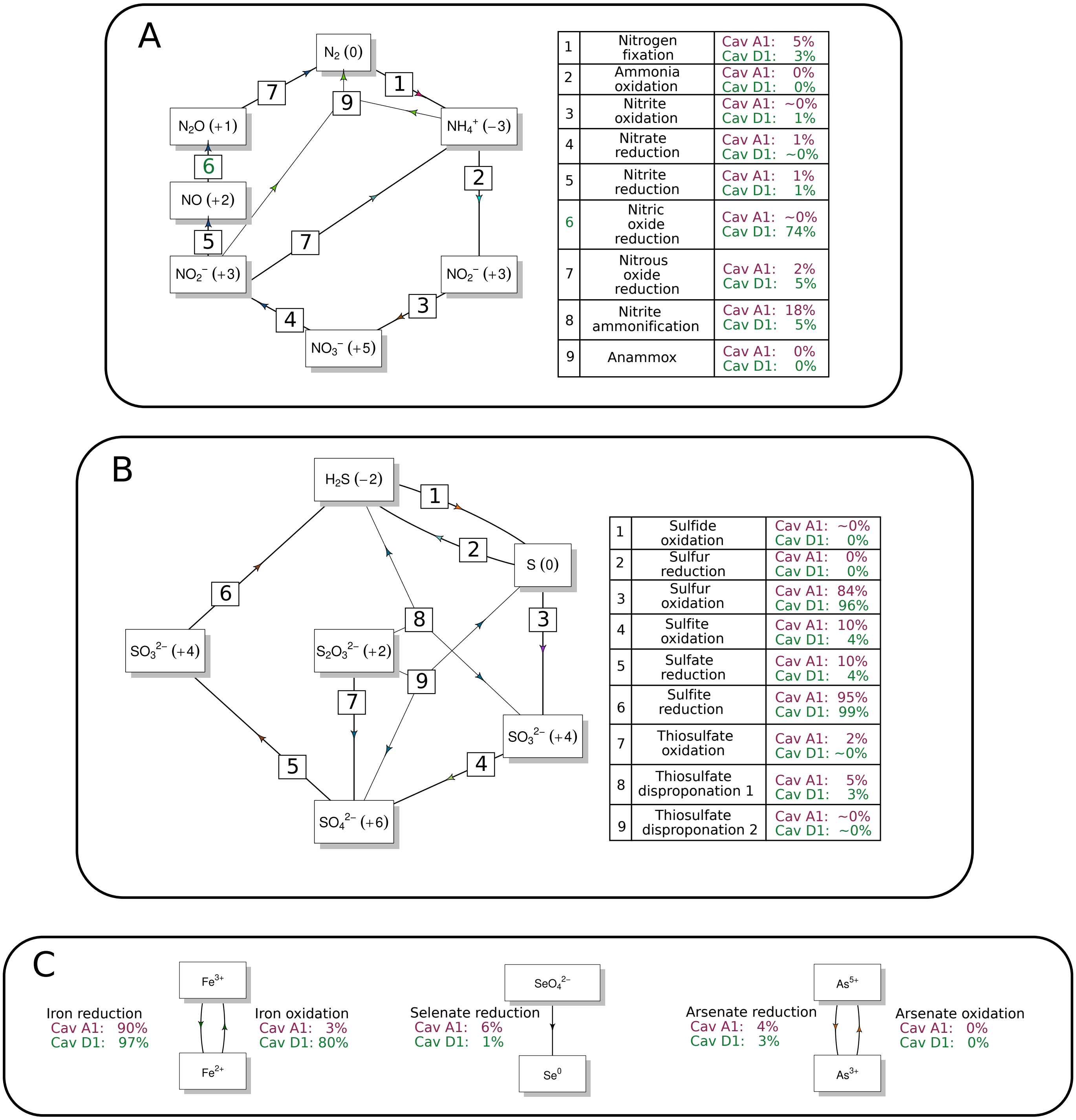


**Figure 3.** Projection of the metabolic functions derived for the Schwab dataset on the nitrogen (A), sulfur (B), and other cycles (C). Like the carbon cycle diagram in Fig. 2B in the main text, the diagrams are taken from [2]. The diagrams are completed with weights of the functions, given by the relative abundances of the microorganisms in the two considered samples of the Schwab dataset (Cav. A1 and Cav. D1).

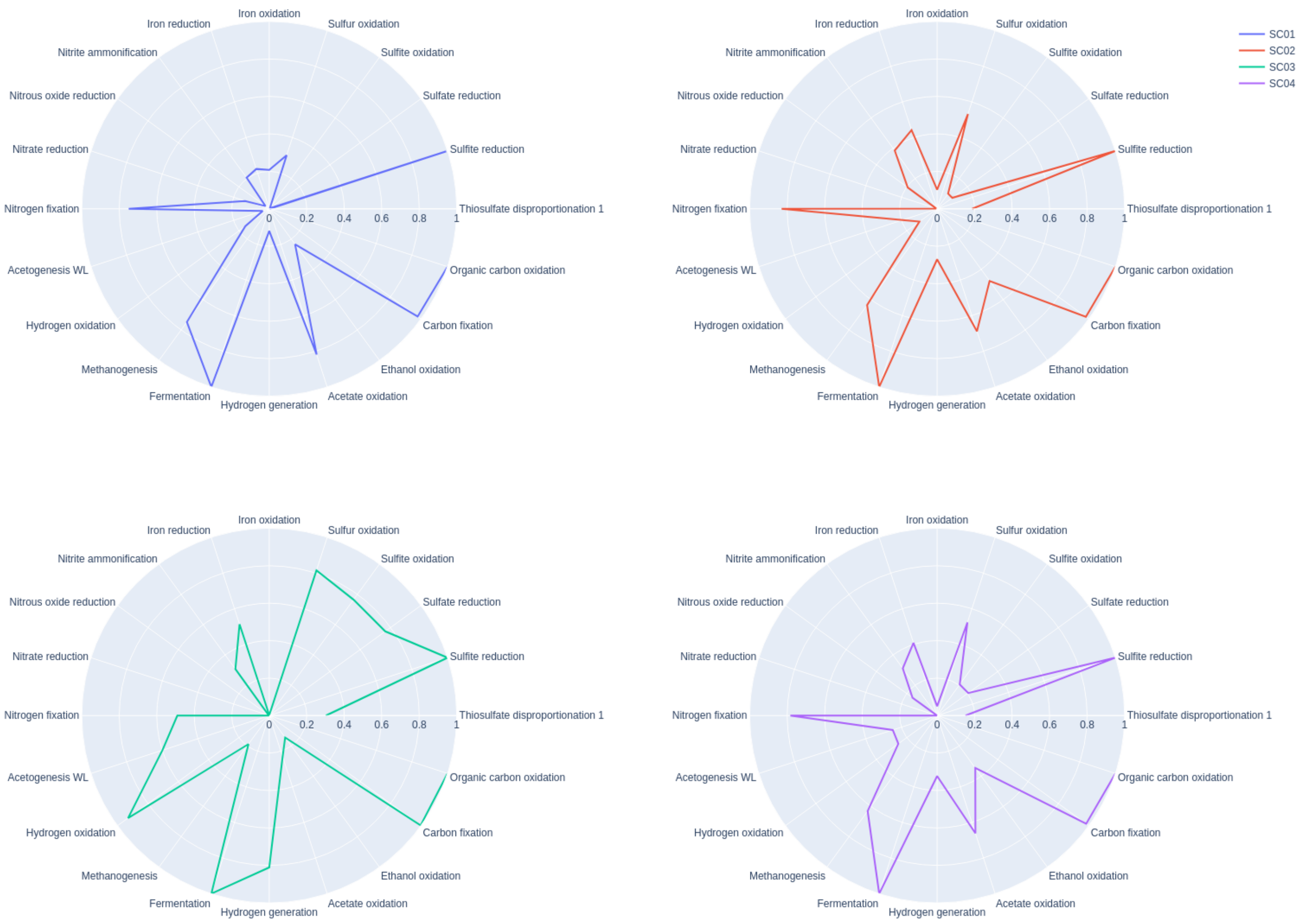


**Figure 4.** Polar plots showing, for each major metabolic function, the relative abundance of microorganisms found in the samples in the Bordenave dataset [3]. Only functions having at least 10% of relative abundance in at least one of the samples are shown.

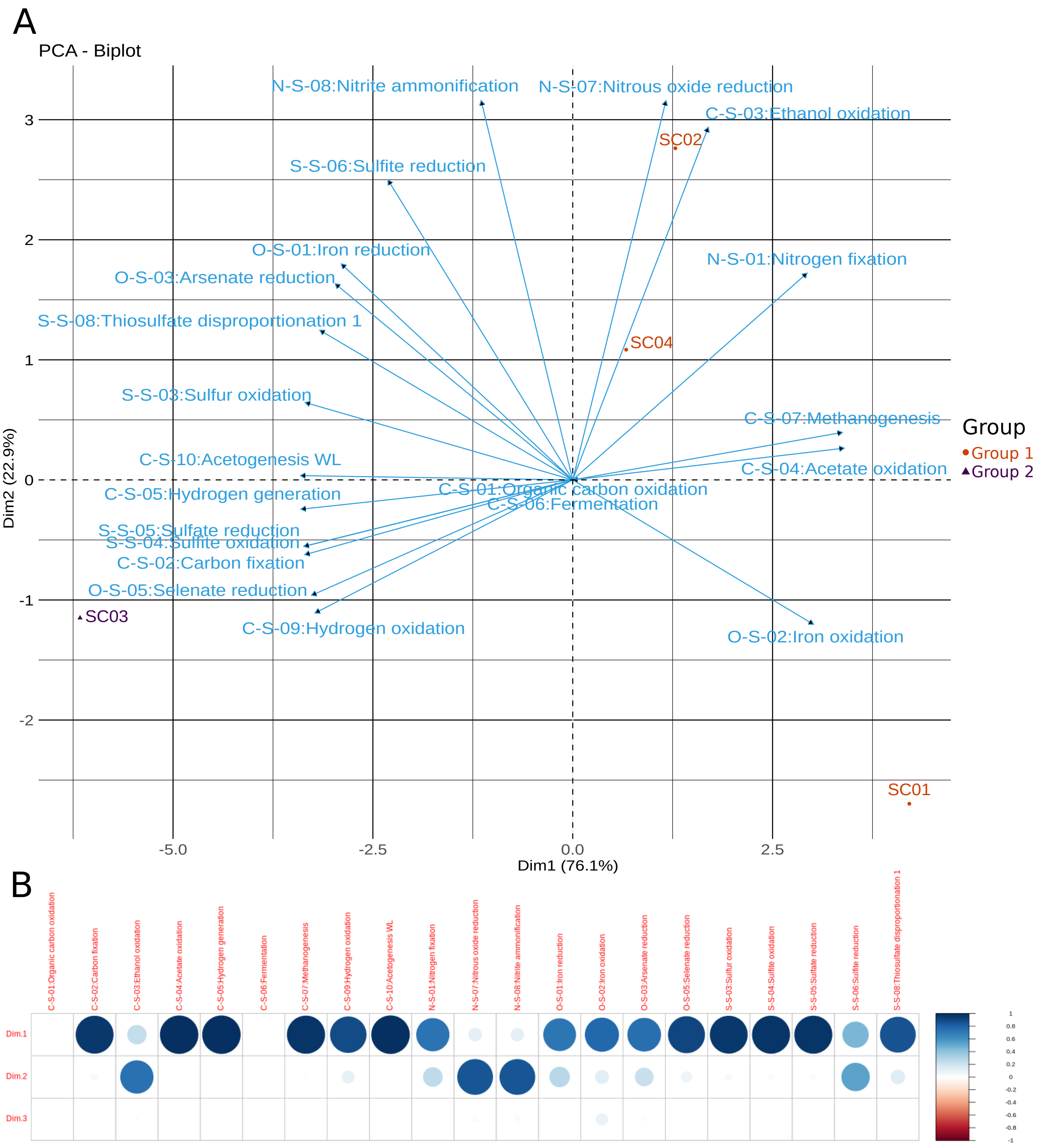


**Figure 5.** Principal Component Analysis (PCA) of the results obtained for the Bordenave dataset. A. Projection on the first two PCA dimensions of the relative abundances of metabolic functions reconstructed from the samples in the Bordenave dataset [3]. The vectors in the biplot represent the metabolic functions and the points the salt caverns. The latter are clustered in two groups located on opposite sides of the origin: one group for SC01, SC02, and SC04, and one group for the remaining cavern SC03. Negative correlations are observed between metabolic functions. For example, the vectors representing acetogenesis and acetate oxidation point in opposite directions. B. Plot of the correlation of the relative abundances of metabolic functions in the Bordenave dataset samples with the 9 PCA dimensions.

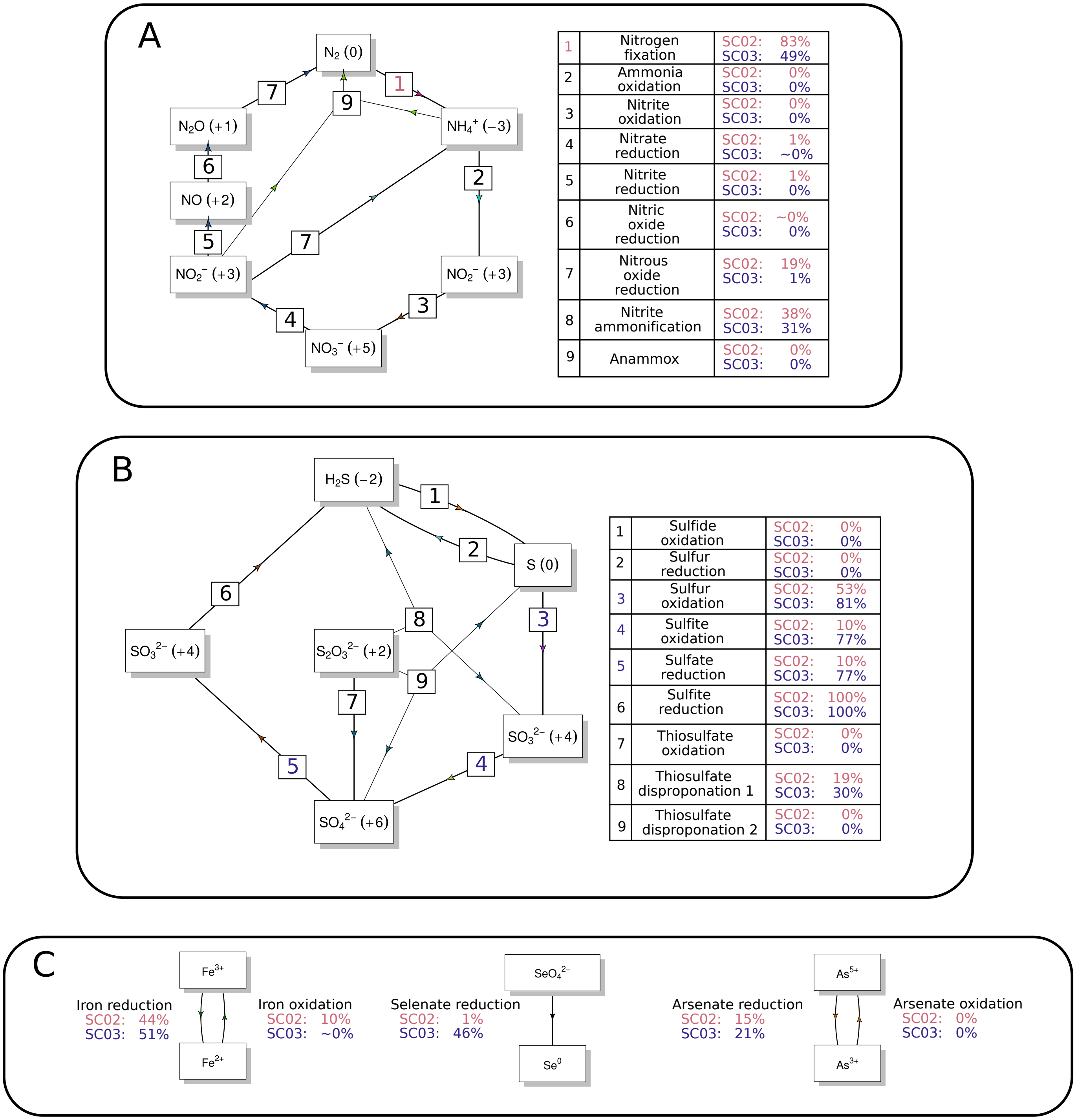


**Figure 6.** Projection of the metabolic functions derived for the Bordenave dataset on the nitrogen (A), sulfur (B), and other cycles (C). Like the carbon cycle diagram in Fig. 2D in the main text, the diagrams are taken from [2]. The diagrams are completed with weights of the functions, given by the relative abundances of the microorganisms in the two considered samples of the Bordenave dataset (SC02 and SC03).

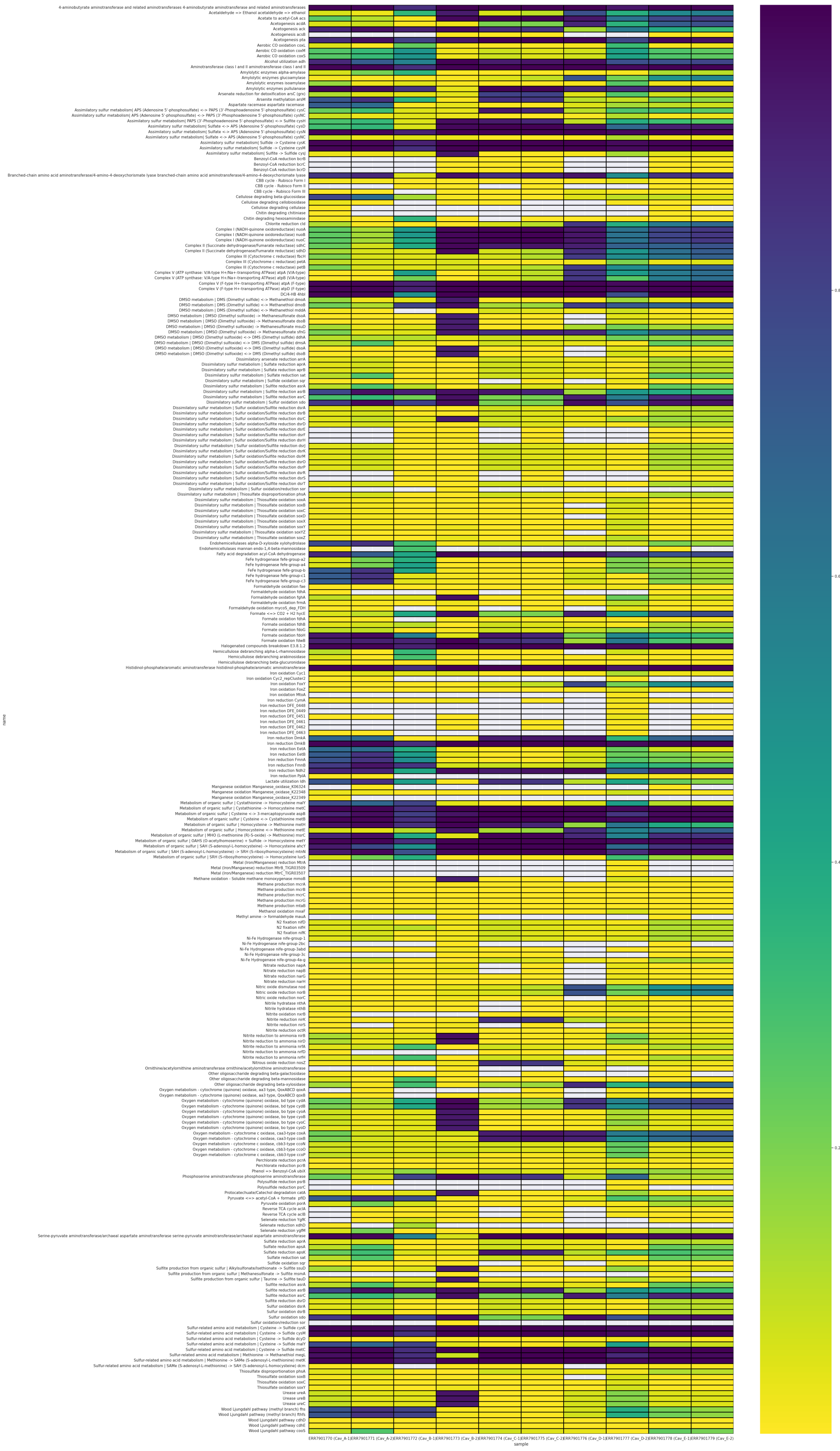

**Figure 7.** Heatmap showing, for each metabolic function, the relative abundance of microorganisms found in the samples in the Schwab dataset.

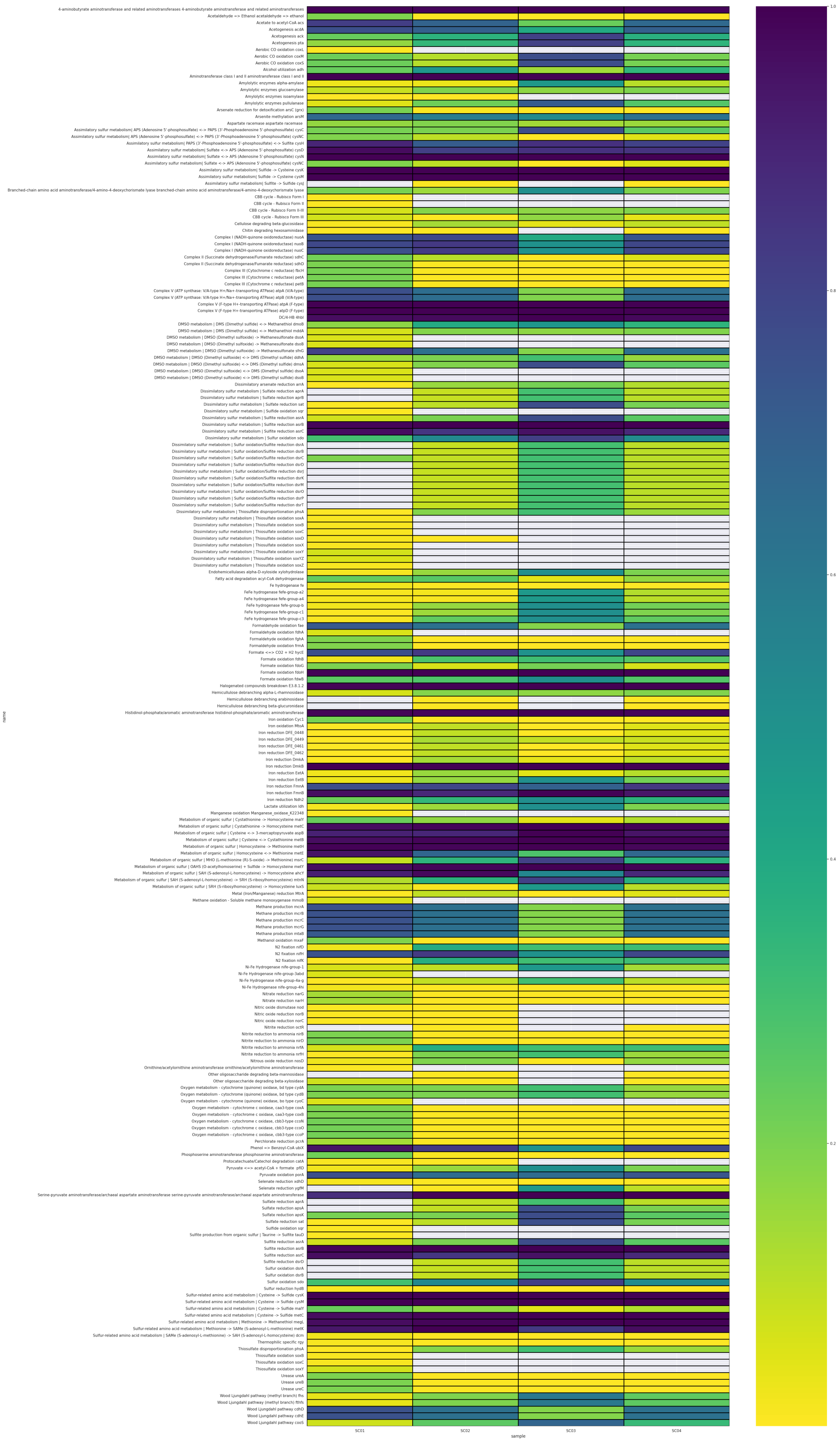


**Figure 8.** Heatmap showing, for each metabolic function, the relative abundance of microorganisms found in the samples in the Bordenave dataset.

### References


[1] Laura Schwab, Denny Popp, Guido Nowack, et al. “Structural analysis of microbiomes from salt caverns used for underground gas storage”. In: *Int J Hydrogen Energy* 47.47 (2022), pp. 20684–94.

[2] Zhichao Zhou, Patricia Q. Tran, Adam M. Breister, et al. “METABOLIC: high-throughput profiling of microbial genomes for functional traits, metabolism, biogeochemistry, and community-scale functional networks”. In: *Microbiome* 10.1 (2022), p. 33.

[3] Sylvain Bordenave, Indranil Chatterjee, and Gerrit Voordouw. “Microbial community structure and microbial activities related to CO2 storage capacities of a salt cavern”. In: *Int Biodeterior Biodegradation* 81 (2013), pp. 82–7.
